# Urbanization and altitude impact on gut microbiome of an Andean frog (*Pristimantis unistrigatus*)

**DOI:** 10.1101/2021.11.10.468176

**Authors:** Elena Catelan Carphio, Diego F. Cisneros-Heredia, Andrés Caicedo, Paúl A. Cárdenas

## Abstract

The analysis of the intestinal microbiome in amphibians provides insights of the anthropogenic environmental impact. *Pristimantis unistrigatus* is an Andean amphibian species whose distribution has been recorded in Ecuador and Colombia, ranging from endemic elfin forests to urban gardens. In this study, we focus on the analysis of the *P. unistrigatus* microbiome 16S rRNA gene sequencing. A total of 32 specimens of *P. unistrigatus* were collected and analyzed from 4 locations in the Valley of Quito, Ecuador, characterized by several urban environments and altitudes. The results show that the relative abundance of bacteria is significantly different amongst groups. Clostridiales are proportionally more abundant in rural and lower altitude locations, while Erysipelotrichaceae, Desulfovibrionaceae, Enterobacteriaceae, Bacteroidaceae and Lachnospiraceae are found at higher elevations. These results highlight the importance of the evolution of the microbiome as a tool of adaptation and survival of amphibians in the present-day changing ecosystems undergoing anthropogenic stresses.

**Importance:** Amphibians constitutes one of the groups most vulnerable to environmental alterations. Due to their sophisticated reproductive and breeding requirements and their permeable skins to breathe, amphibians are compulsively studied as ecological indicators. The destruction of pristine habitats occurred all over the planet in recent decades has caused a catastrophic decline in amphibian populations for many species everywhere. Ecuador, being one of the most biodiverse country on Earth, hosts a huge variety of amphibians, thus offering a unique possibility of studying the biology of the amphibian species living in its ecosystems, and how they adapt to changing habitats. A direct way to diagnose the status of an amphibian population is to study the gut microbiome of the individual specimens. The gut microbiome is closely related to the host’s health and so to its ability to adapt and survive. An important output of this study is to offer indications and tools useful to conservation programs before irreversible damages are caused to the habitats and the amphibians’ populations still thriving in them.

## Introduction

Microorganisms, primarily bacteria, but fungi and viruses as well, have developed complex relationships with bigger host organisms, such as mammals, fishes, amphibians, arthropods, among others. Recent studies have shown that bacteria play an essential role in disease resistance, health conditions and adaptation of their host to biotic and abiotic stressors (Jimenez & Sommer, 2017; Colston & Jackson, 2016; Chang et al., 2016; Bahrndorff et al., 2016), the relationship between the microbial community and its host is considered mutualistic, commensal, symbiotic and even pathogenic (Jimenez & Sommer, 2017; Karl et al., 2018). Every living being has a countless number of bacterial cells in their organism, but regardless of the specific number, it’s their interaction as complex communities and the influence on their host what makes the understanding of their role so important (Colston & Jackson, 2016). Research development in this area has allowed a deeper comprehension of the microbial diversity in any sample of interest, primarily by the sequencing of fragments of the bacterial 16S rRNA gene. The characterization of microbial communities has turned out to be faster and affordable thanks to the development of next generation sequencing and other modern technologies (Colston & Jackson, 2016; Bahrndorff et al., 2016; Jimenez & Sommer, 2017).

Since the Human Microbiome Project (HMP) started in 2008, the majority of gut microbiome studies have concentrated in mammals, particularly humans, though prompting a number of questions about how the gut microbiome functions in animals other than mammals. The animal microbiome has evolved with their hosts, being shaped by their genotype, life stage and the ecological and physiological conditions (Bahrndorff et al., 2016). In turn, the microbiome is functional to the host’s nutrient acquisition, immune response (Mashoof et al., 2013; Colombo et al., 2015), behavior (Banas et al., 1988), development (Knutie et al., 2017a; Warne et al, 2017; Chai et al. 2018), reproduction and, most importantly, host’s health condition (Colston & Jackson, 2016; Chang et al., 2016; Warne et al, 2017; Pereira et al., 2017). Any alteration of the microbiome composition may hamper its normal capabilities, turning the host more vulnerable to the effects of unfavorable environmental conditions (Jimenez & Sommer, 2017; Zhang et al., 2016; Mu et al., 2018; Kohl et al., 2014; Huang et al. 2018).

The gut microbiome develops a symbiotic relationship with the host’s organism, a fundamental stage aimed at capturing nutrients (Sugita et al. 1984; Chang et al., 2016; Warne et al, 2017), fermenting fibers and synthetizing essential amino acids (Kohl et al., 2014). The importance of these relationship has been just recently explored to attempt an explanation of the ecological success of certain species in their environment (Kohl et al., 2014; Warne et al, 2017; Huang et al., 2018), and recent studies have shown its implications as relevant for wildlife conservation (Jimenez & Sommer, 2017; Bahrndorff et al., 2016). However, despite the expansion of this area of research, most of the information comes from data collected from mammals. The number of studies on amphibians is almost negligible and notwithstanding, the majority of these studies are from animals kept in confinement (Warne et al, 2017; Benno et al., 1992), thus leading to information quasi useless for the purposes of wild animal conservation (Colston & Jackson, 2016), since the number of environmental disruptors (Knutie et al., 2017b) that are presumably unaccounted and/or systematically neglected when toads are kept in captivity (Smith & Stoskopf, 2007).

The increase of human encroachment in pristine ecosystems has caused changes in species abundance and distribution, especially amphibians (Pounds et al., 1999). The amphibian population has declined since the 1980s all over the world, the tropical Pacific Ocean region being the most affected, including countries such as Ecuador. Anthropogenic stressing processes have dramatically affected amphibians’ and other vertebrate species’ ecosystems, causing amongst several effects an increase of temperature. (Pounds, 2001). Temperature and moisture alterations are the two phenomena in climate change that have directly affected the amphibian biology (Carey & Alexander, 2003), altering body physiology, thereby affecting the gut microbiome (Amato et al. 2013).

During the period between 1990 and 2016, solely 21 studies of microbiome in wildlife frogs have been carried out and published, and only 5 of them used NGS (Colston & Jackson, 2016). Furthermore, none of the mentioned studies reports data from Ecuador, a region belonging to a biodiversity hotspot of amphibians’ species. Specifically, Ecuador is the fourth most diverse country in terms of amphibians’ species in the world, hosting 609 species, 558 of them of the Anura order, the *Pristimantis* genus being by far the most abundant one (BioWeb, 2019). *Pristimantis unistrigatus* is a common species inside this group, a relatively small-size frog well adapted to the habitats within the 2200-3400 m Andean valleys, from southern Colombia to central Ecuador. Across this elevation range, vegetation changes from forested regions in sub-temperate wet and humid temperate regimes to pastures, ditches, shrubs, crops, forest edges and urban areas. *P. unistrigatus* is a terrestrial nocturnal species thriving in leaf litter, tolerant to human intrusion and therefore commonly found even in urban areas and farmlands. A number of characteristics has allowed this species to be encountered in different ecosystems stressed by various degrees of human encroachment and disruption, to the point that its IUCN conservation status is LC. Reasonably, the gut microbiome is playing a crucial role in the evolution and adaptation of *P. unistrigatus,* allowing the potential to overcome environmental threatening processes like habitat fragmentation, acid deposition and the emergence of new diseases, parasites and predators. To support these arguments, in this study we focus to determine whether the composition of the microbial communities within *P. unistrigatus* species has significantly changed across 4 niches with different altitude and degree of anthropogenic impact, thus enhancing its ecological resilience and ability of adaptation and coping with novel habitats and environmental recesses.

## Material and Methods

### Sample collection

Forty specimens (10 from each of four sites) of *P. unistrigatus* male frogs were collected within the Metropolitan District of Quito, namely from two rural areas : Hacienda Sierra Alisos (0.408628W 78.589187S, 3200 m above sea level (asl), HAP) and Hacienda San Francisco (0.441422W 78.561351S, 2830 m asl, HCM); and from two urban areas: Urbanización San Martin (0.177726W 78.496875S, 2880 m asl, SMQ) and Urbanización La Colina (0.317348W 78.434793S, 2510 m asl, VH), between January and May, 2018 (Figure 1). HAP, HCM, SMQ and VH as their respective identifying acronyms throughout this study.

**Figure 1:**
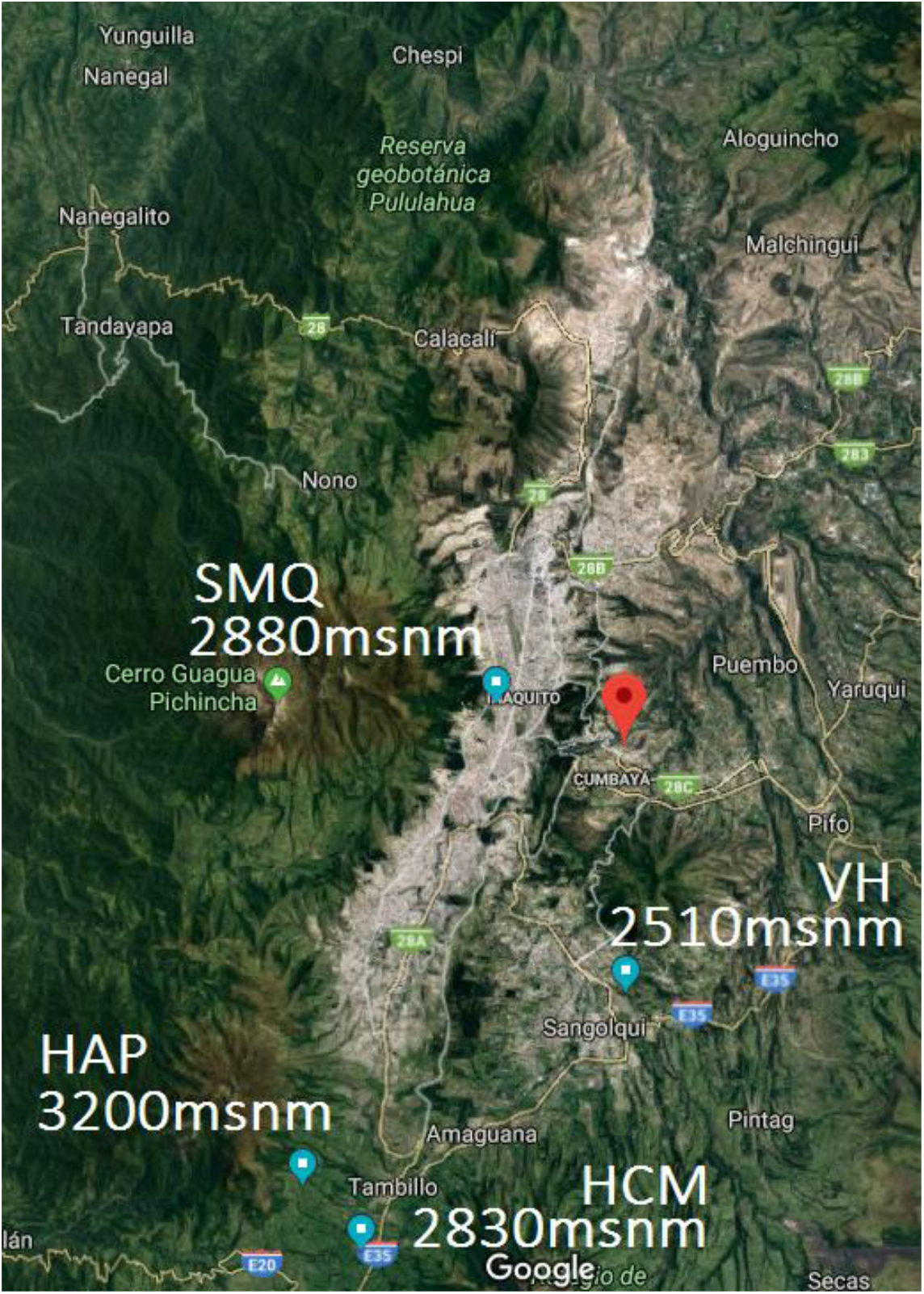
Map of each location in Quito

Due to its nocturnal habits, the collecting and sampling activities were carried out during the night hours. To attract females, *P. unistrigatus* males use a characteristic wooing chant, that well served to guide us in order to locate them in the pitch-darkness. After spotting a male individual, just prior capture, the body temperature was measured using a remote infrared thermometer sensor. To keep pristine its microbiological composition, the specimen was carefully manipulated by one of us during the capture operation, after dressing surgical gloves. Afterward, each collection spot was labeled using a distinguishing marker, and few observables like soil and substrate composition, local environmental temperature and humidity, and wind speed were measured. Prior the final sacrifice, the frog’s body was thoroughly cleansed with distilled water for 30 seconds, both in ventral and dorsal position. Then, a sample of the abdomen was taken with the help of a sterile swab, and a further dorsal sample was collected with a second sterile swab; both samples were finally stored inside 5 ml sterile Falcon tubes.

Each specimen was sacrificed on the spot. The frog was placed inside an Erlenmeyer device connected to an 8 kq CO2 tank. The CO2 flowed through the sealed Erlenmeyer for 10 seconds, and then the connecting valve was closed. The frog was kept inside the Erlenmeyer during further 10 to 30 seconds, until fainted. Then the individual was extracted from the Erlenmeyer and its reflexes tested. With the help of a sterile swab, it was checked whether the blinking reflex was absent; similarly, there had to be no reflex of recovering the ventral position while resting on the back.

Finally, the extraction of the tissues could be carried out. For the dissection, the frog was dorsally positioned, and its abdomen opened with a H-shaped cut. The entire digestive system was skillfully extracted, then the stomach and intestine were separated and stored into two different sterile 5 ml Falcon tubes. Eventually, that same night, the samples were transported and brought to our laboratory at San Francisco de Quito University (USFQ), to be preserved within a freezer at −80 °C temperature.

### Diet analysis

The stomach sample was defrosted at the USFQ Zoology Laboratory. Its entire content was extracted, every found prey was classified at order level with the help of a stereomicroscope (Hyslop, 1980). The index of relative importance (IRI) was calculated to quantify its dietary significance. We measured transversal and longitudinal typical dimensions of each prey applying the formula IRI= O% (N% + V%), where O%, N%, V% are the percentage of occurrence, relative abundance, and measured volume of each prey category, as found within every stomach, respectively (Chang et al., 2016). The volume of a prey was calculated applying the formula (Dias & Rocha, 2007):

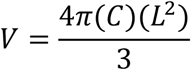

where C is half of the length and L is half of the width. The estimate of the food and spatial niche breadth between habitats is given by the Simpson’s index of diversity (B) (Dias & Rocha, 2007):

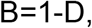

where

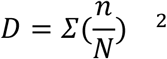

Here *n* is the total number of organisms of a given species, while N is the total number of organisms of all species. Thus, the allowed range of D is between 0 and 1, where 0 corresponds to no diversity, and 1 to the maximum degree of diversity (Dias & Rocha, 2007). The overlap possibly existing amongst the four locations was quantified by the coefficient of symmetry of overlapping, given by:

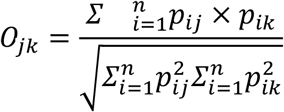

Again, its allowed range is between 0 and 1, 0 corresponds to no overlap, while 1 describes complete overlap (Dias & Rocha, 2007). The number of prey items, relative abundance and prey volume were analyzed with MiniTab 18 version with a T-test to determine differences between urban and rural habitats, and higher and lower habitats. **Intestinal microbiome analysis**

We analyzed the intestinal microbiome of 32 frogs (8 from each of the four locations), a subset of the 40 specimens we collected, because of budget limitations and DNA concentration constraints. Intestinal microbial DNA was extracted using PureLink Microbiome DNA Purification Kit (Stool Samples). For each sample, we amplified the V3 and V4 hypervariable 16S rRNA region using the primer set Bakt_341F: CCTACGGGNGGCWGCAG and Bakt_805R: GACTACHVGGGTATCTAATCC. The sequencing was carried out on a Miseq 300bp PE at Macrogen Inc. (Song et al., 2018). The bioinformatic analysis was performed using Qiime2 2018.4 version, every sequence was assigned to a barcode, demultiplexing the sequences and determining how many sequences were obtained per sample and their quality. For the denoising and control processes of the sequences, DADA2 was used to detect and correct the sequenced data (Mashoof et al., 2013).

The Alpha diversity analyses that we carried out amounted to Shannon’s diversity index (Medina et al., 2017), Observed OTUs (Chai et al., 2018), Faith’s Phylogenetic Diversity (Bletz et al., 2016) and Pielou’s Evenness (Knutie et al., 2017b), and these metrics were computed for each frog using Qiime2. As for the Beta diversity, we calculated Unweighted and Weighted UniFrac distances between samples using Qiime2 (Knutie et al., 2017a). The sequencing depth for the analysis amounted to 2823 sequences. This value was chosen based on the number of sequences in the LBI012 sample. The visualizations resulting from the prior analysis were generated with the Emperor tool (Chai et al., 2018).

### Availability of data and materials

The 16S rDNA sequences identified in this study have been deposited in the European Nucleotide Archive under the accession numbers from ERS3017357 to ERS3017387. Frog samples were stored at the Zoology Laboratory of the USFQ. The swaps of skin microbiome were stored at the Microbiology Institute of the USFQ.

## Results

### Diet differentiation amongst habitats

#### Urban vs. Rural

A total of 32 *P. unistrigatus* individuals, from which 17 came from rural habitats (HAP and HCM) and 15 from urban habitats (VH and SMQ) were analyzed. A total of 133 individual prey items were identified to 11 orders (Table 1). There is a clear difference in the presence of certain groups of prey between urban and rural habitats. The higher index of relative importance (IRI) score in the Rural habitats is for the order Aranae (IRI=88680,719), followed by Diptera (IRI=66746,739). While in the Urban habitats the most abundant order was Coleoptera (IRI=63126,431), followed by Isopoda (IRI=44150,5414) (Table 1, Figure 2A). The number of prey items (p=0,896), relative abundance (p=1,00) and prey volume (p=0,722) were not significantly different between the four locations (Supplementary Table 1). But according to the Simpson’s Index of Diversity the urban habitats (B=0,818) were a little more diverse than the rural habitats (B=0,729) and the overlapping between the urban and rural habitats was O=0,864 (Supplementary Table 2). Lepidoptera, Neuroptera, Dermaptera, Isopoda and Harpacticoida were only present in the urban habitats (Table 1).

**Table 1:**
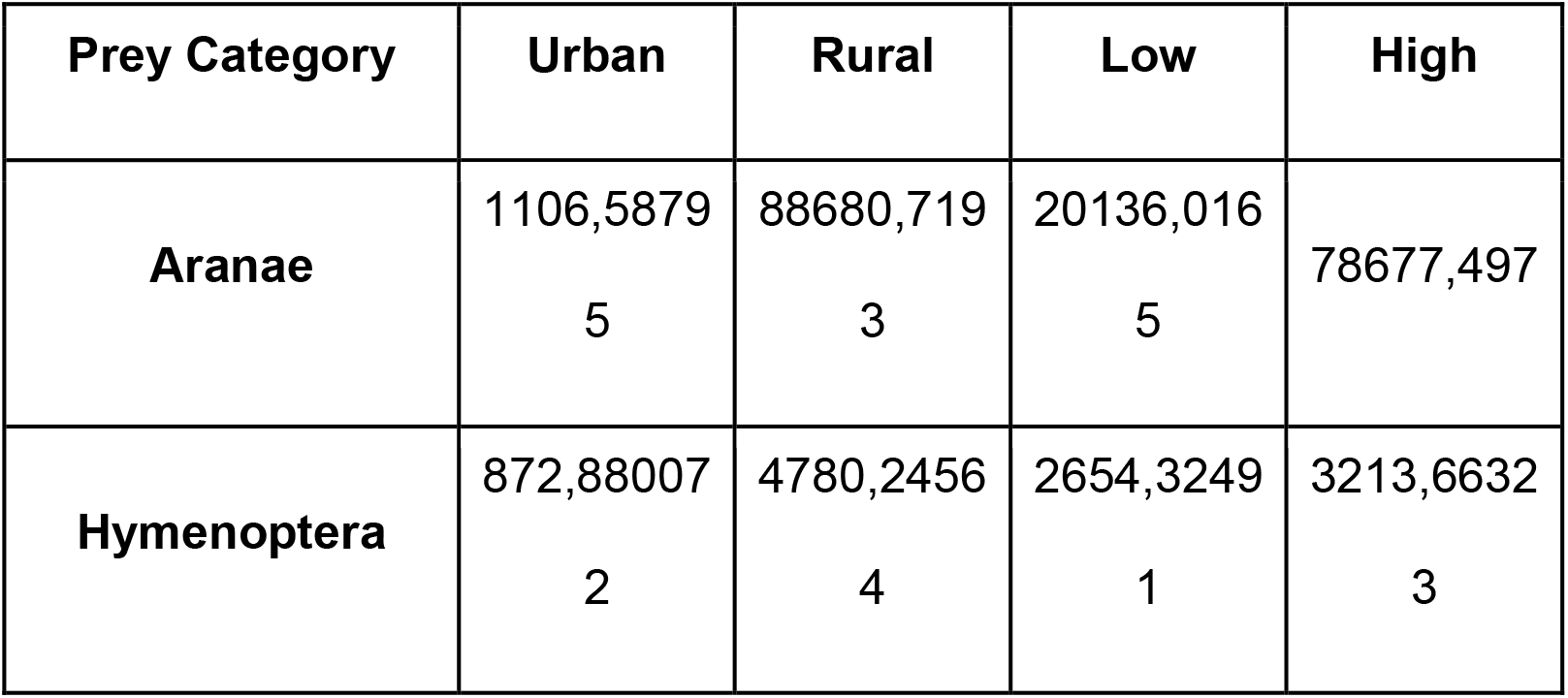
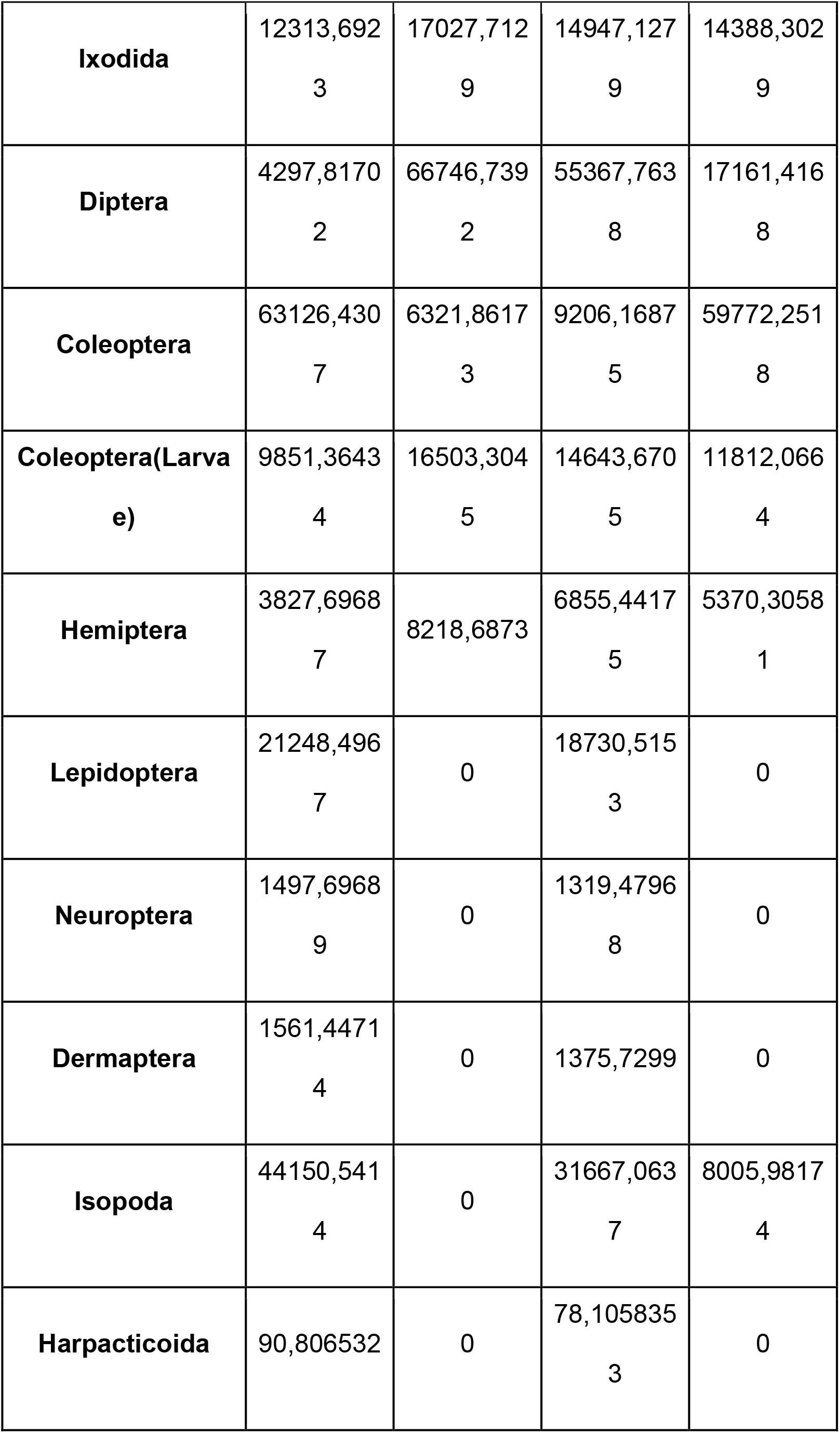
Index of relative Importance of the Stomach contents of *Pristimantis unistrigatus* in the four sites of collection

**Figure 2:**
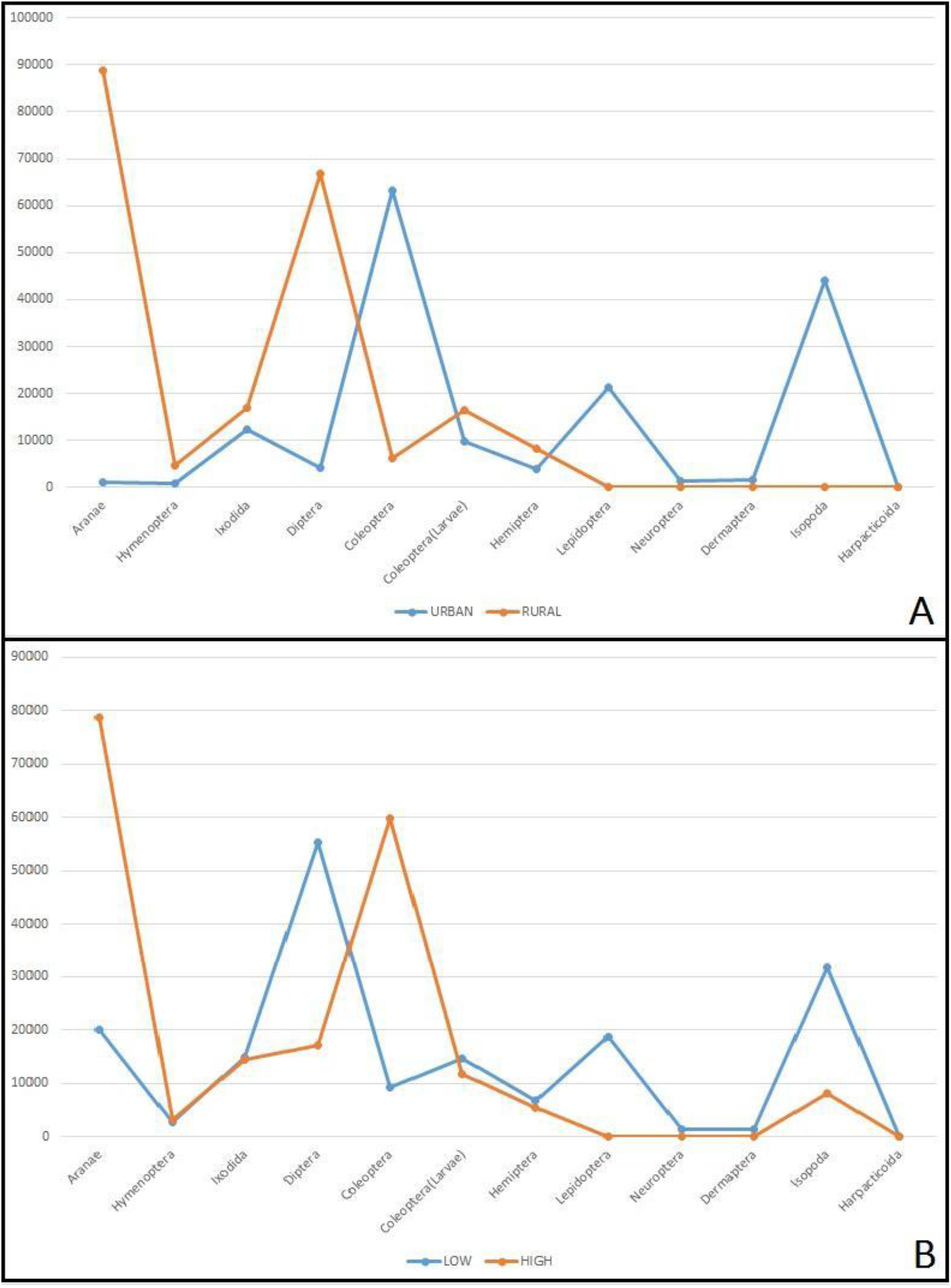
Differences in Index of Relative Importance of frog’s preys between urban and rural (A) and between low and high (B) habitats.

#### High vs. Low altitude

A total of 32 *P. unistrigatus*, from which 15 came from the higher locations (HAP and SMQ) and 17 from the lower locations (HCM and VH) were analyzed. A clear difference in the preference of certain preys between the high and low altitude habitats is observed in Figure 2B. In the higher habitats the order Aranae has the higher index of relative importance (IRI) score (IRI=78677,497), followed by Coleoptera (IRI=59772,251). While in the lower habitats Diptera has the higher IRI score (IRI=55367,763), followed by Isopoda (IRI=31667,063) (Table 1, Figure 2B). The number of prey items (p=0,400), relative abundance (p=1,00) and prey volume (p=0,840) were not significantly different between the higher and lower habitats (Supplementary Table 1). But according to the Simpson’s Index of Diversity the lower habitats (B=0,803) were by little more diverse than the higher habitats (B=0,751) and the overlapping between the higher and lower habitats was almost complete with O=0,951. Lepidoptera, Neuroptera, Dermaptera and Harpacticoida were only present in the lower habitats (Table 1).

### Composition of the intestinal microbiome

Gut bacterial communities were investigated using 16S rRNA gene in 31 individuals of *P. unistrigatus* at Quito. The sequencing resulted in a total of 364 243 high-quality sequences from all the samples. Multiple rarefaction curves generated from the observed OTUs reached a plateau phase suggesting that high sampling coverage was achieved in all samples (Figure 3). From the taxonomic analysis a diverse community structure was revealed, dominated by members of the phyla Firmicutes (mostly Clostridia), Proteobacteria (mostly Gammaproteobacteria) and Bacteroidetes (mostly Bacteroidia) (Figure 4). The concentration of certain groups of bacteria was different between the four locations, suggesting a different microbiome composition among the four populations of Andean frogs.

**Figure 3:**
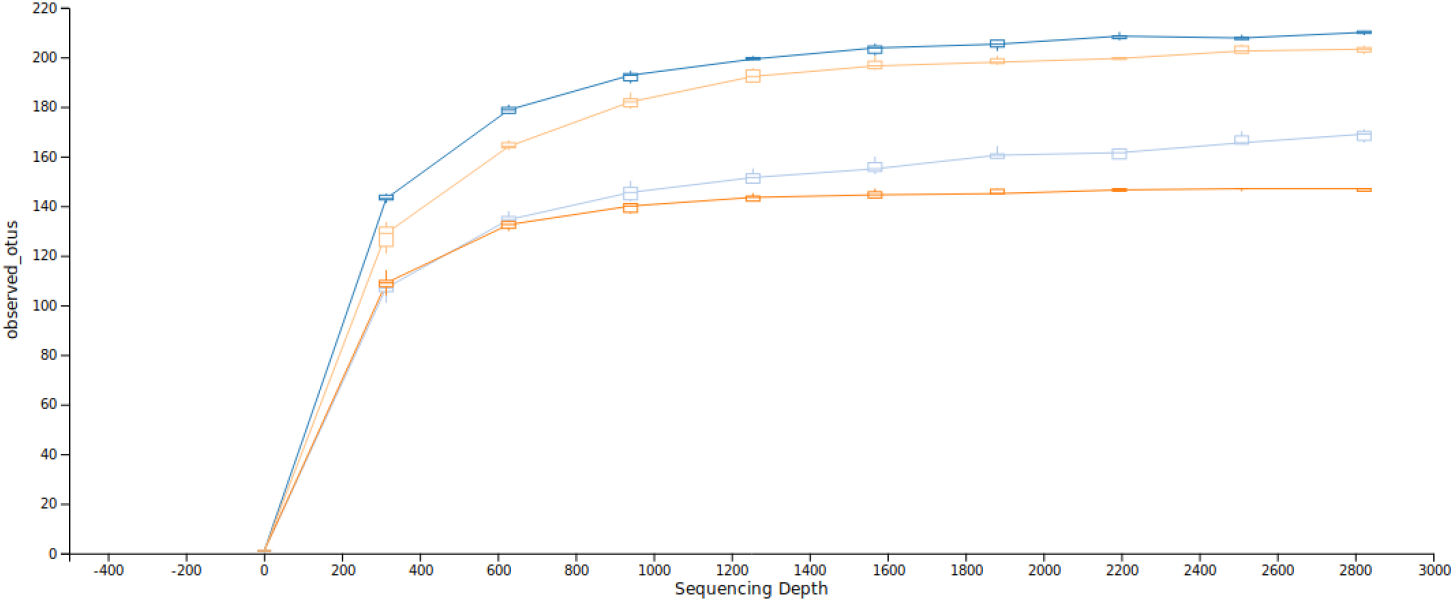
Rarefaction curves of Andean frogs based on Ilumina MiSeq sequencing. Horizontal axis number of sequenced data and vertical axis the observer number of the operational taxonomic units.

**Figure 4:**
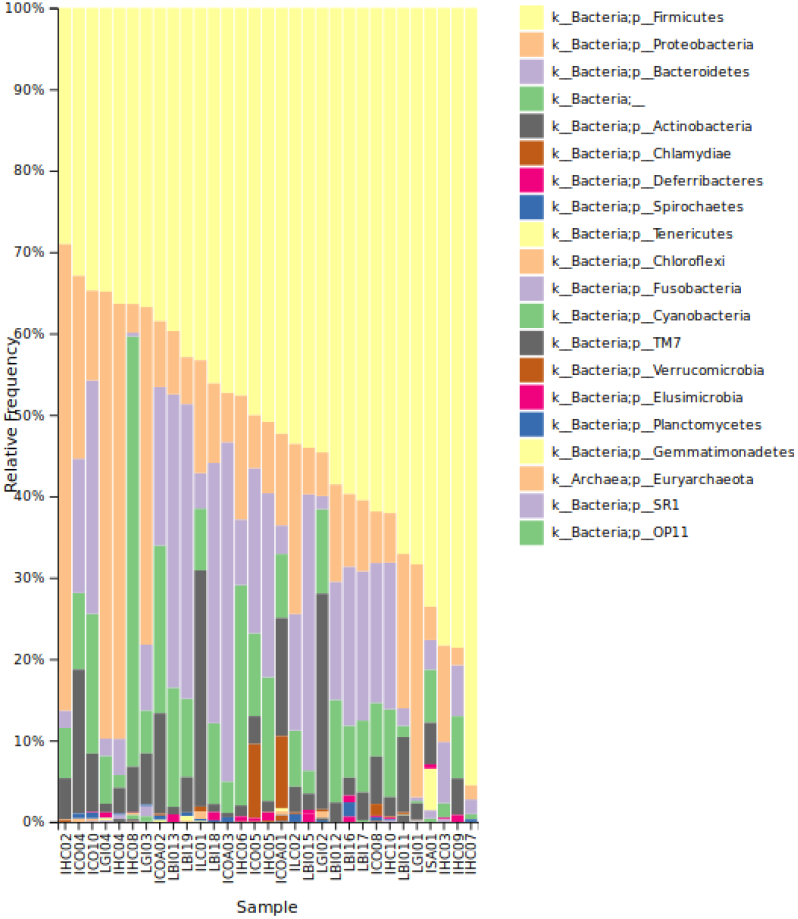
Relative frequency of microbial taxa based on Ilumina MiSeq.

The alpha diversity (faith and evenness) of the intestinal microbiome was estimated by Kruskal-Wallis. The faith between all groups was not significant (*P*=0.499), as consequence the richness of species in every location was not different and the pairwise analysis showed no significant *P-*values in any of the groups (Supplementary Table 3). Nevertheless, the evenness of the intestinal microbial community was different between all groups (*P*=0.009), showing the species of bacteria inside each group were different among them (Supplementary Table 3). The pairwise analysis showed significant differences between HAP-HCM (*P*=0.012), HAP-SMQ (*P*=0.015), and HCM-VH (*P*=0.022); but there were no differences between HAP-VH (*P=*0.643), HCM-SMQ (*P=*0.100) and SMQ-VH (*P=*0.084) (Supplementary Table 3).

The beta diversity measures the diversity of species between the intestinal microbial communities of each group. Based on weighted UniFrac distance with PCoA analysis (Figure 5A) there isn’t a defined cluster for any of the groups but there is a tendency for each individual of the group to be close to each other, there were significant differences in the pairwise PERMANOVA results with 999 permutations between HAP-HCM (*P=*0.036), HAP-SMQ (*P=*0.010), HCM-VH (*P=*0.008) and SMQ-VH (*P=*0.006), while between HAP-VH (*P=*0.173) and HCM-SMQ (*P=*0.140) there were not significant differences (Supplementary Table 4). On the other hand, the PCoA analysis based on the unweighted UniFrac distance (Figure 5B) showed no define clusters within the groups, in the pairwise PERMANOVA results there were significant differences between HAP-HCM (*P=*0.044), HAP-SMQ (*P=*0.012), HAP-VH (*P=*0.006), HCM-VH (*P=*0.004) and SMQ-VH (*P=*0.012), while there were not significant differences between HCM-SMQ (*P=*0.619) (Supplementary Table 4). The PCoA analysis based on the Jaccard distance shows 2 clusters (HAP and VH), while the other two groups were dispersed through the entire graphic (Figure 5C).

**Figure 5:**
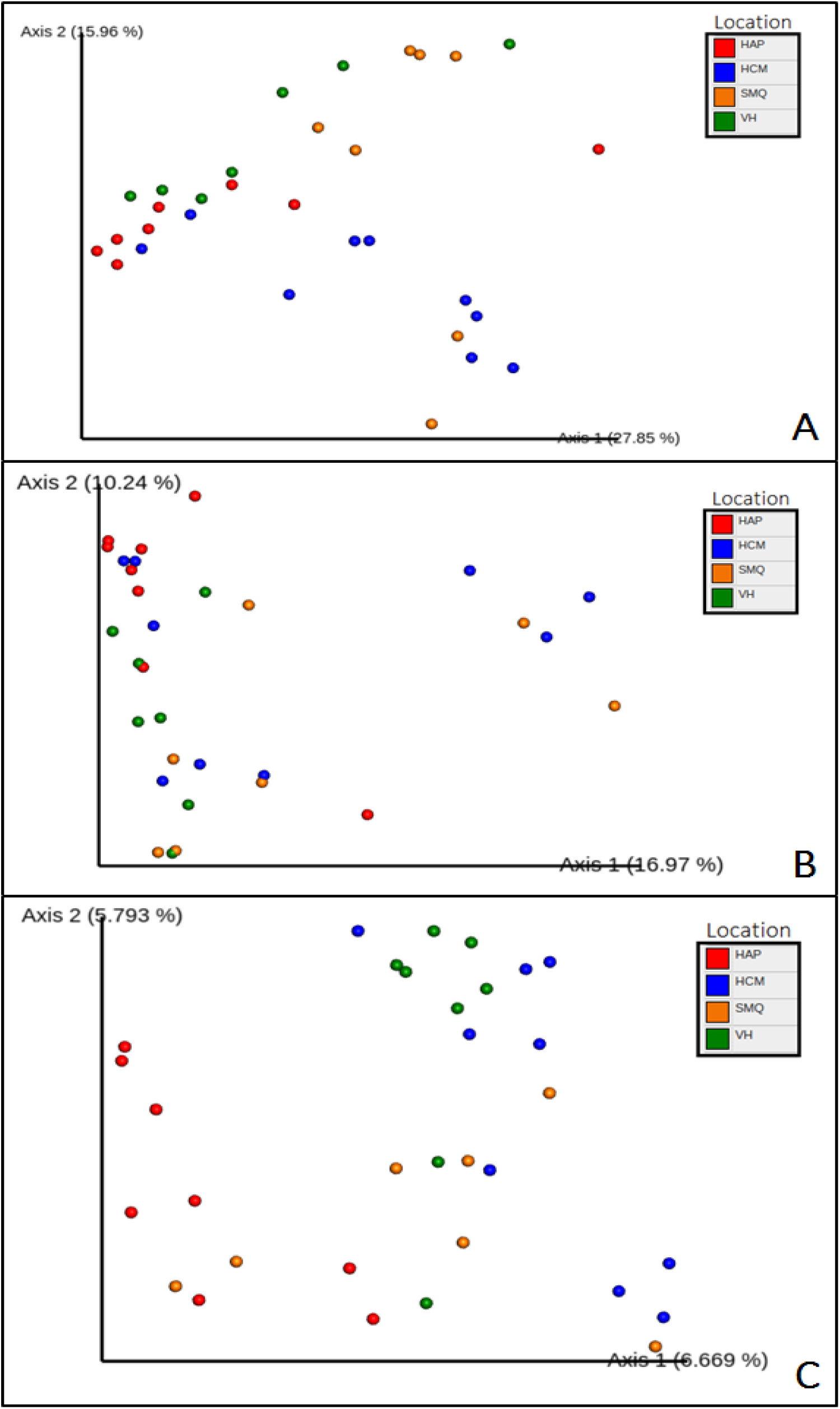
Beta diversity of intestinal microbial communities among the four elevations based on the weighted UniFrac distance (A), unweighted UniFrac distance (B) and Jaccard Scores (C)

Differential abundance analysis using balances in gneiss revealed that proportions of *Clostridiales* were a lot higher in the lower and rural locations (Figure 6). Bacteroidaceae, Erysipelotrichaceae, Desulfovibrionaceae, Enterobacteriaceae, Bacteroidaceae and Lachnospiraceae were more present in the higher locations (Figure 6A). Lachnospiraceae, Ruminococcaceae and Erysipelotrichaceae were more present in urban locations (Figure 6B).

**Figure 6:**
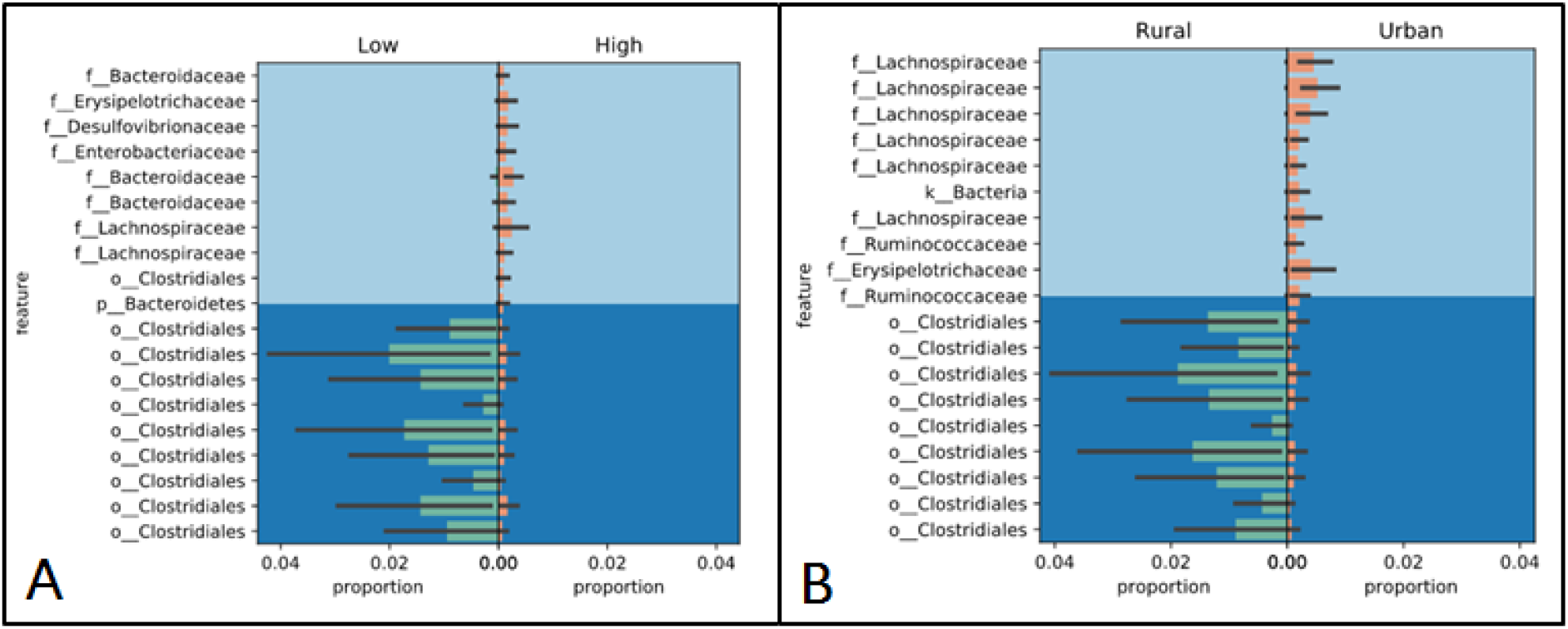
Proportion Plot of differential abundance analysis using balances in gneiss according to the elevation (A) and urbanization (B).

#### Variables of importance

##### Soil temperature

In PERMANOVA results based on the weighted UniFrac distance (permutations=999) showed significant differences between VH and SMQ at the range of temperatures 14-15°C vs. 15-16°C (*P=*0.047) (Supplementary Table 5) while on the other pair of ranges there were no significant differences. Between VH and HCM at the range of temperatures 13-14°C vs. 14-15°C (*P=*0.032) and 14-15°C vs. 15-16°C (*P=*0.030) (Supplementary Table 5), while on the other pair of ranges there were no significant differences. Additionally, comparisons between all the other groups there were no significant differences among the ranges of temperatures.

On the other hand, in the PERMANOVA results based on the unweighted UniFrac distance (permutations=999) between VH and HCM showed significant differences at the range of temperatures 14-15°C vs. 15-16°C (*P=*0.029) (Supplementary Table 6), while on the other pair of ranges there were no significant differences. Between SMQ and HCM at the range of temperatures 12-13°C vs. 15-16°C (*P=*0.036), and 14-15°C vs. 15-16°C (*P=*0.040) (Supplementary Table 6), while on the other pair of ranges there were no significant differences. Between all the other groups there were no significant differences among the ranges of temperatures.

In all the pair of locations the soil temperature seemed to be a factor of influence in the composition of the gut microbiome, but the PCoA analysis (Supplementary Figure 1) showed a correlation that every location had a specific soil temperature, so taking all of this into consideration the soil temperature was a variable of confusion but did not restructure the microbial community.

##### Relative environmental humidity

The PERMANOVA results based on the weighted UniFrac distance (permutations=999) showed significant differences between SMQ and HAP at the humidity ranges 90-93% vs. 94-97% (*P=*0.025) and 90-93% vs. 98-100% (*P=*0.029) (Supplementary Table 7), while on the other pair of ranges there were no significant differences. Between HAP and HCM at the humidity range 90-93% vs. 98-100% (*P=*0.019) (Supplementary Table 7). Between all the other groups there were no significant differences among the humidity ranges. Moreover, the PERMANOVA results based on the unweighted UniFrac distance (permutations=999) showed significant differences between SMQ and HAP at the humidity ranges 90-93% vs. 94-97% (*P=*0.037) and 90-93% vs. 98-100% (*P=*0.020) (Supplementary Table 8), while on the other pair of ranges there were no significant differences. Between all the other groups there were no significant differences among the humidity ranges.

#### Location of importance

Differential abundance analysis using balances in gneiss revealed that proportions of *Clostridiales* were a lot higher in HCM in comparison to SMQ, while Lachnospiraceae, Eubacteriaceae, Phyllobacteriaceae and Intrasporangiaceae were more abundant in SMQ than in HCM (Supplementary Figure 1). Even though *Bacteroides*, and *Clostridium*, are present in HAP and VH, the proportion in which certain species are present in one of the two locations is completely different (Supplementary Figure 2). *Parabacteroides* and Veillonellaceae are more present in VH, while Ruminococcaceae is more present in HAP (Supplementary Figure 2).

## Discussion

In the present study, we used four locations to determine if the level of urbanization and the altitude would be a factor that had a potential role in changes in the structure of the gut microbiome of *P. unistrigatus*. The factors influencing the association of microbial communities inside a host are the primary interest in microbial ecology, which can be diet, physiological conditions and the host species (Sugita et al., 1985). This host-microbiome symbiosis may facilitate the survival rate of a species facing the rapid ecosystemic changes around the globe (Bletz et al., 2016). The dominance of certain groups of bacteria can be beneficial, while the presence of other groups can be harmful for the frog (Karl et al., 2018). Is so that the study of the composition of the intestinal microbiome of bigger organisms may be an insight of the state and evolutionary story of a certain population.

Diet structure and changes can be one of the external factors that alter the composition of the intestinal microbiome. Is so that frogs’ gut microbiomes can be influenced by soil microorganisms due to the ingestion of the preys covered in soil bacteria found in their actual niche (Huang et al., 2018). The lack of differences in stomach contents of Andean frogs between urban vs. rural and high vs. low habitats may reflect that. However even with the human impacts there is not a significant disturbance to alter the faunal species composition. The diet analysis showed that the food volume, number of prey items and relative abundance of prey categories were not significantly different between the four locations.

The environment is changing, and a lot of habitats are being destroyed, so prey resources also change (Chang et al. 2016), this is reflected in different prevalence of the preys according to the origin of the sample. Indeed, Aranae had the higher IRI score in the rural and high altitude habitats, which could suggest a preference from frogs towards this group. While Diptera (Rural and Low altitude habitats) and Coleoptera (Urban and High-altitude habitats) have completely contrasting habitat preferences, and so we see a higher abundance of these groups in these ecosystems. Although further research is needed to understand the causes that lead to these differences. The diverse degree of human impact among all the groups suggested that the diet would have changed between the individuals. But the absence of statistical differences and the high overlap of species between the groups showed that the dietary tendency of the frogs was the same among all groups, meaning this couldn’t be a factor of influence to the differences in the composition of the gut microbiome.

The structure of the gut microbial communities are highly determined by environmental factors of the niche where the host develops (Chang et al. 2016). The adaptation to certain ecosystems can change the host’s tolerance, behavior and interaction with its surroundings, which can alter the gut microbiome (Huang et al., 2018). The amphibians’ intestinal microbiome is similar to that of mammals and birds. And like that it can vary due to external factors (Benno et al., 1992). The intestinal microbiome analysis showed a clear variation throughout the various locations, suggesting the environmental factors within the urbanization and altitude affected the microbial communities. The various variables measured in each group did not showed a direct influence in the microbial communities, due to the lack of significant *P-*values. Excluding, the relative environmental humidity that in certain cases showed significant differences between the ranges of humidity. The metabolism of amphibians requires high quantities of water, when the levels of humidity changes can cause a level of stress in the organisms (Silva et al., 2012). And so several responses start to regulate stress, which can mediate the growth of the gut microbiome and elucidate their activity. Intestinal cells can produce neuroendocrine hormones that directly affect the microbial communities (Karl et al., 2018).

The most abundant groups of bacteria in the intestinal microbiome of *P. unistrigatus* were Firmicutes (mostly *Clostridiales*), Proteobacteria and Bacteroidetes. Different studies around the globe have shown the same dominance of these groups in the intestinal microbiome of other species of anurans. However, looking closer the bacterial composition patterns were specific to each species (Vences et al., 2016; Weng, et al., 2016). A question for further investigation remains open: which is the role of each group inside the intestinal microbiome. For example, Proteobacteria which was one of the most abundant groups throughout all the samples, could help with the metabolism of amino acid substrates even in low concentrations (Beebee & Wong, 1992). On the other hand, predictions by dynamic interaction modeling between Firmicutes and Proteobacteria have suggested interactions among these microorganisms, but the complexity of this microbiome has made it difficult to have experimental data to examine these relationships (Weng et al., 2017).

The dominance of Clostridia in the low and rural groups suggest there are anaerobic conditions in the gastrointestinal system of *P. unistrigatus* (and could be one of the major differences with the skin microbiome) (Vences et al., 2016). Still this group is also related to inflammatory responses and reducing the microbiome diversity in the human intestine (Karl et al., 2018). In addition, changes like this in the organization of the microbiome could lead to alterations to the immune system being the primary sensor for microbes and their metabolites (Weng et al., 2016). The development and function of the innate and adaptive immune system of toads is highly influenced by the colonization of different microorganisms (Weng et al., 2016). An increase in environmental stress can lead to a corruption of the intestinal microbiome which also make anurans susceptible to bacterial infections leading to septicemia and even death of the organisms (Fedewa, 2006).

Stress can lead to dysbiosis in the gastrointestinal system harming the host health (Karl et al., 2018). For example, in hibernating frogs the metabolism decreases, reducing also the functioning of the intestinal microbiome. Revealing also that this could change the species interaction and functioning with the ecosystem (Weng et. al., 2016). Studies have shown that the presence of filamentous bacteria can prevent the colonization of pathogenic bacteria in the gastrointestinal system of anurans. Then again, the presence of filamentous bacteria in the intestine can vary due to diet and external variables (Klaseen et al., 1993).

Amphibians have evolved an intimate relationship with their microbial communities (Boni & Battaglini, 1964). Gut microorganisms can reflect an evolutionary selection driven by the external environment (Huang et al., 2018). The structure of microbial communities differs between elevations. High altitude can be an environmental stressor for the functioning of the gastrointestinal system. It can lead to the loss of appetite, nausea, abdominal pain, among others in humans. Although intestinal epithelial cells work normally under a gradient of oxygen, the lower proportion of oxygen has changed the composition of the intestinal microbiome (Karl et al., 2018). This suggests that the environmental conditions at high and low location does contribute to the composition of the gut microbiome. Different studies showed that there were relationships of the diversity and richness of bacteria with altitude patterns, moreover in Andean regions there were not previous reports in soil bacteria influenced by it (Medina et al., 2017). Also, a study of the human gut microbiome composition through an altitude range has shown structural differences among the groups. Where Firmicutes were more abundant in the group with the higher altitude, and Bacteroidetes was more abundant in the group of lower altitude (Li & Zhao, 2015), which is inverse to the results in this research where Firmicutes were more abundant in the lower groups.

Although there is no consensus on what a ‘healthy’ microbiome looks like, there is a level of agreement on the characteristics where the composition is favorable for success of a certain species (Karl et al., 2018). The understanding of how these micro-ecosystems work can generate a new focus of study of the ecology of amphibians around the world and so, the understanding of how the communities of this species in Ecuador could be adapting to the anthropogenic impacts in their ecological niches. The response and adaptation of the intestinal microbiome over environmental stressors can be promoting or degrading the health of the host. Is that so that the intestinal microbiome could be a factor that helps the host to outcome short- and long-term changes in their environment (Karl et al., 2018). Being this one of the first researches of frogs’ intestinal microbiome in Ecuador, further investigation is needed to truly understand the dynamics inside and between the microbiome, the host and the environment where this species develops.

## Acknowledgments

We thank Benjamin Arias and Mateo Flores for their help in the specimen collection. Lucia Fiallos and Daniela Pazmiño for their help in the extraction and preparation of samples for the DNA sequencing. And finally, Belen Prado for the assistance in the use and interpretation of the Qiime2 results.

## Appendix

### Supplementary Figures

**Supplementary Figure 1:**
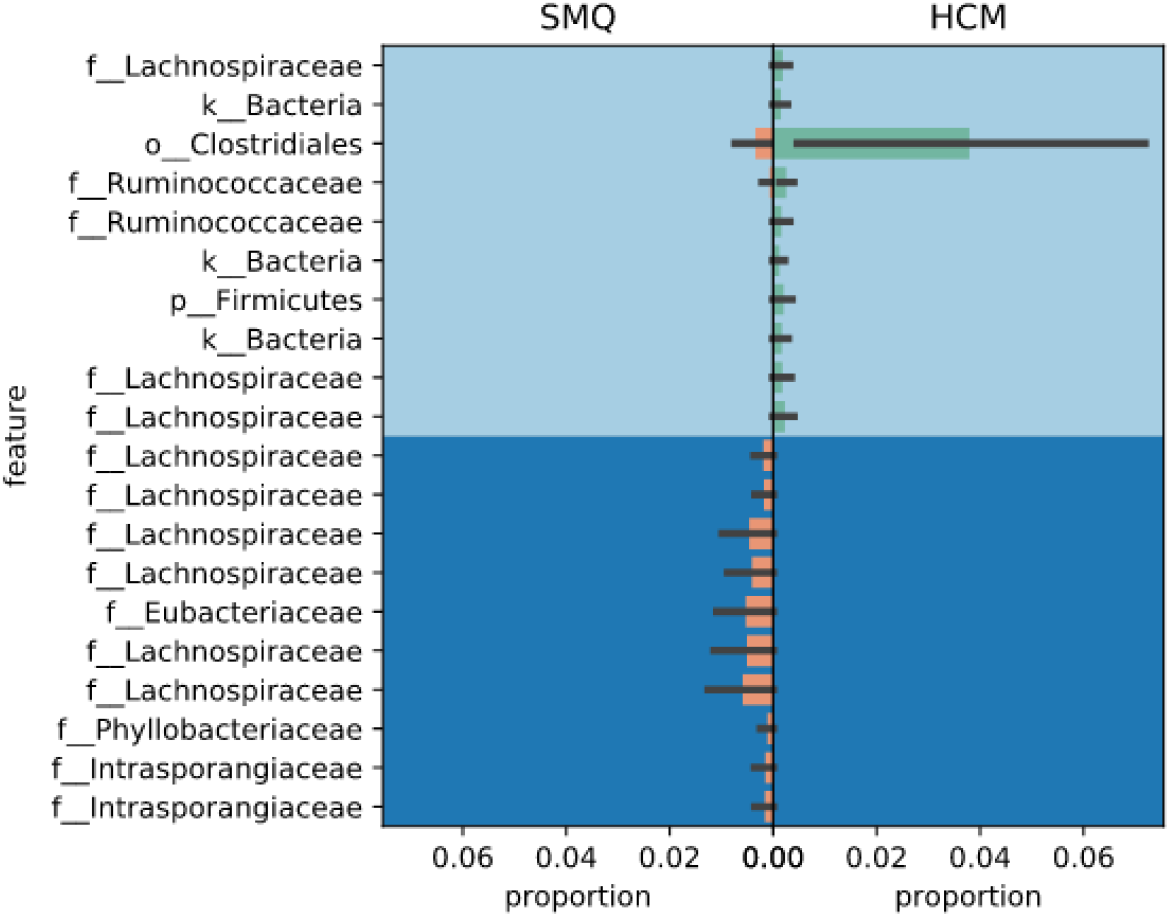
Proportion Plot of differential abundance analysis using balances in gneiss comparing two locations.

**Supplementary Figure 2:**
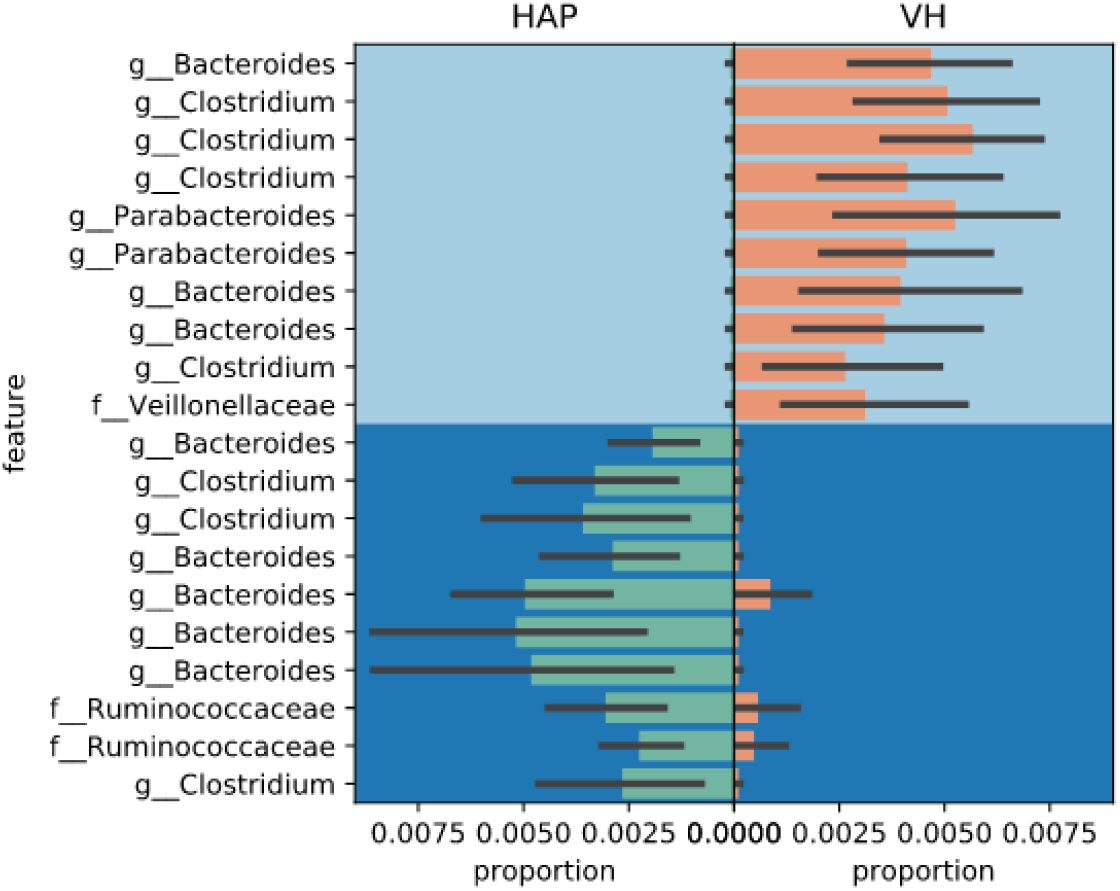
Proportion Plot of differential abundance analysis using balances in gneiss comparing two locations.

### Supplementary Tables

**Supplementary Table 1:** T-test of stomach contents of *P. unistrigatus* in the four sites of collections

|  | **Urban vs.**<br>**Rural** | **Low vs.**<br>**High** |
| --- | --- | --- |
| **Number of Prey Item** | 0.896 | 0.400 |
| **Relative abundance** | 1.000 | 1.000 |
| **Volume of Prey**<br>**(mm<sup>3</sup>)** | 0.722 | 0.840 |

**Supplementary Table 2:** Simpson’s index and Coefficient symmetry of overlapping in the four sites of collections

|  | **Simpson's**<br>**Index** | **C. S. of**<br>**Overlapping** |
| --- | --- | --- |
| High | 0.751249519 |  |
| Low | 0.803688281 |  |
| High vs. Low |  | 0.951151658 |
| Rural | 0.729258559 |  |
| Urban | 0.818359375 |  |
| Urban vs.<br>Rural |  | 0.86388799 |

**Supplementary Table 3:** *P-*Values of Alpha diversity analysis (Evenness and Faith) between the four sites of collection

| Location | Evenness | Faith |
| --- | --- | --- |
| HAP vs. HCM | 0.012 | 0.500 |
| HAP vs. SMQ | 0.015 | 0.132 |
| HAP vs. VH | 0.643 | 0.643 |
| HCM vs. SMQ | 0.100 | 0.368 |
| HCM vs. VH | 0.022 | 0.873 |
| SMQ vs. VH | 0.084 | 0.337 |

**Supplementary Table 4:** *P-*Values of Beta diversity analysis based on weighted and unweighted UniFrac distances between the four sites of collection

| Location | Unweighted | Weighted |
| --- | --- | --- |
| HAP vs. HCM | 0.044 | 0.036 |
| HAP vs. SMQ | 0.012 | 0.010 |
| HAP vs. VH | 0.006 | 0.173 |
| HCM vs. SMQ | 0.619 | 0.140 |
| HCM vs. VH | 0.004 | 0.008 |
| SMQ vs. VH | 0.012 | 0.006 |

**Supplementary Table 5:** *P-*Values of Beta diversity analysis based on weighted UniFrac distances between the sites of collection comparing the ranges of soil temperature

| Group 1 | Group 2 | VH vs. SMQ | VH vs. HCM |
| --- | --- | --- | --- |
| 12-13°C | 13-14°C | 0.098 | - |
|  | 14-15°C | 0.262 | - |
|  | 15-16°C | 0.274 | - |
|  | 16-17°C | 1.000 | - |
| 13-14°C | 14-15°C | 0.521 | 0.032 |
|  | 15-16°C | 0.426 | 0.134 |
|  | 16-17°C | 0.246 | 0.058 |
| 14-15°C | 15-16°C | 0.047 | 0.030 |
|  | 16-17°C | 0.428 | 0.067 |
| 15-16°C | 16-17°C | 1.000 | 0.681 |

**Supplementary Table 6:** *P-*Values of Beta diversity analysis based on unweighted UniFrac distances between the sites of collection comparing the ranges of soil temperature

| Group 1 | Group 2 | VH vs. HCM | SMQ vs. HCM |
| --- | --- | --- | --- |
| 12-13°C | 14-15°C | - | 0.663 |
|  | 15-16°C | - | 0.036 |
|  | 16-17°C | - | 0.595 |
| 13-14°C | 14-15°C | 0.064 | - |
|  | 15-16°C | 0.104 | - |
|  | 16-17°C | 0.089 | - |
| 14-15°C | 15-16°C | 0.029 | 0.040 |
|  | 16-17°C | 0.385 | 0.391 |
| 15-16°C | 16-17°C | 0.461 | 0.386 |

**Supplementary Table 7:**
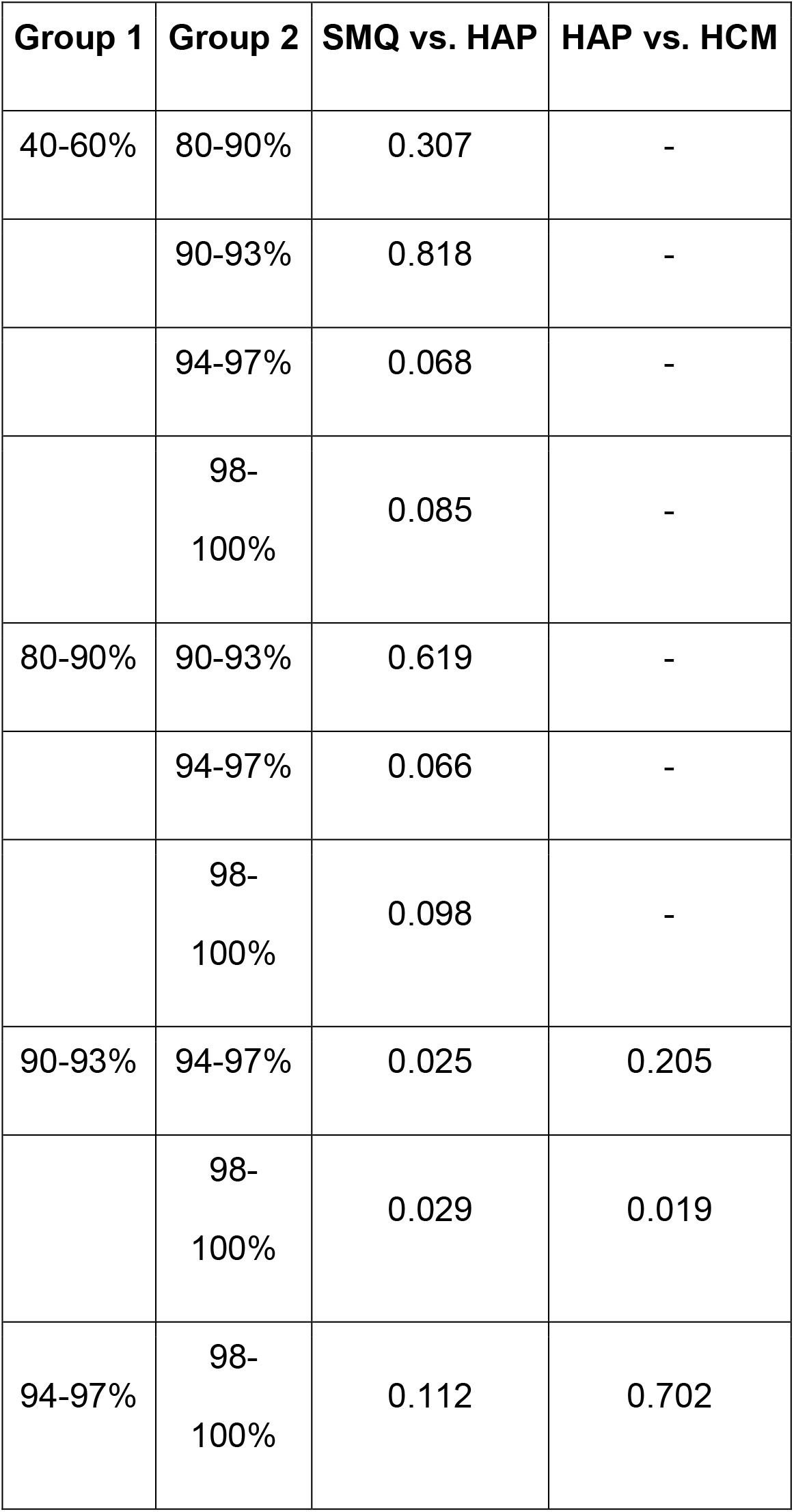
*P-*Values of Beta diversity analysis based on weighted UniFrac distances between the sites of collection comparing the ranges of environmental relative humidity

| Group 1 | Group 2 | SMQ vs. HAP | HAP vs. HCM |
| --- | --- | --- | --- |
| 40-60% | 80-90% | 0.307 | - |
|  | 90-93% | 0.818 | - |
|  | 94-97% | 0.068 | - |
|  | 98-100% | 0.085 | - |
| 80-90% | 90-93% | 0.619 | - |
|  | 94-97% | 0.066 | - |
|  | 98-100% | 0.098 | - |
| 90-93% | 94-97% | 0.025 | 0.205 |
|  | 98-100% | 0.029 | 0.019 |
| 94-97% | 98-100% | 0.112 | 0.702 |

**Supplementary Table 8:** *P-*Values of Beta diversity analysis based on unweighted UniFrac distances between the sites of collection comparing the ranges of environmental relative humidity

| Group 1 | Group 2 | SMQ vs. HAP |
| --- | --- | --- |
| 40-60% | 80-90% | 0.0331 |
|  | 90-93% | 0.674 |
|  | 94-97% | 0.066 |
|  | 98-100% | 0.109 |
| 80-90% | 90-93% | 0.729 |
|  | 94-97% | 0.060 |
|  | 98-100% | 0.117 |
| 90-93% | 94-97% | 0.037 |
|  | 98-100% | 0.020 |
| 94-97% | 98-100% | 0.054 |

